# Cytotoxic effect of *Rotheca serrata* on cancer cell lines MCF-7 and Neuroblastoma SH-SY5Y

**DOI:** 10.1101/2022.07.01.498410

**Authors:** Jayashree Pandurang Gadade, Swaroopa Amit Patil

**Affiliations:** Shivaji University, Kolhapur, India; Shivaji University, Kolhapur

**Keywords:** *Rotheca serrata*, Neuroblastoma SH-SY5Y, MCF-7 cell line, anticancer activity, MTT assay, viability of cells

## Abstract

Rotheca serrata (Lamiaceae), a highly medicinal plant is used as an antidote for snakebite and the plant possesses medicinal properties like hepatoprotective, antitussive, antioxidant, anticancer, neuro-protective, used in rheumatoid arthritis and is also a α-glucoside inhibitor. This work aimed to study the anticancerous effect of *Rotheca serrata* (root and leaf) on cancer cell lines MCF-7 (breast cancer cell line) and Neuroblastoma SH-SY5Y. The results indicated that the Methanolic extract of *Rotheca serrata* (root and leaf) showed high anticancer activity. Different concentrations of plant extracts (25, 50, 100, 200, 400 μg/ml) were used to study the anticancerous activity, amongst which the significant results were obtained for 400 μg/ml concentration (both root & leaf). Effective anticancer activity against MCF – 7 breast cancer cells was shown in methanoilc extracts and were expressed as IC 50 values; in root (IC 50 value= 61.8259 ± 7.428 μg/ml) and in leaf (IC 50 value = 78.1497 ± 6.316μg/ml). The MTT assay in case of neuroblastoma (SH-SY5Y) cell lines revealed that 400μg/ml concentration of leaf methanolic extract showed effective inhibition of cancer cells with IC 50 value 37.8462 ± 2.957 μg/ml as compared to IC 50 value of root methanolic extract which was 57.0895 ± 2.351 μg/ml.

## Introduction

Day-to-day life has become faster and technical. Many people suffer from various health problems due to changed life styles which has made life hectic. Synthetic drugs have lots of side effects on human health that’s why everybody is trying for natural remedy to cure health problems which have less or no side effects on health. Plants are the best natural source of many drugs. Medicinal plant contains various secondary metabolites which are extracted and used to treat many life-threatening diseases such as heart disease, cancer and skin diseases.

Noncommunicable diseases (NCDs) account for 71% of total deaths worldwide (Mathur, 2020). In India, NCDs were estimated to account for 63% of total deaths cause due to various diseases and cancer was one the leading cause (9%). The higher proportion of cancers associated with use of tobacco was in the North Eastern states, followed by registries in the West and Central regions of India.

Globally, nearly 1 in 6 deaths are is due to cancer [1]. The cancer-causing agents may be internal or external. The internal factors include mutations that occurs during metabolism, immune conditions, mutations which are inherited and hormones. While the external factors which can cause cancer are tobacco, chemicals, radiation and infectious organisms. Cancer is a significant worldwide health problem generally due to the lack of widespread and comprehensive early detection methods, the associated poor prognosis of patients diagnosed in later stages of the disease and its increasing incidence on a global scale [2].

Cancer is the second leading cause of death globally after cardiovascular disease and has been sole reason for nearly accounting for nearly 10 million deaths in 2020 [3]. Breast cancer (19 PBCRs) and cervical cancer (7 PBCRs) were the most common cancers found in women [4]. The highest burden of breast cancer was observed in metropolitan cities. A steady increase in breast cancer in most of the PBCRs including newer PBCRs, poses a great health challenge to women in India [5].

Neuroblastoma, one of the human malignancies reveal spontaneous regression converting an undifferentiated state to complete cellular benign state. This originates from the sympathetic nervous system and is the most frequently occurring extracranial malignancy in childhood [6,7]. Though there are many advances in the diagnostic procedures and standard interventions in the past three decades, neuroblastoma has remained a frightening challenge to clinical and basic scientists [8]. Drug discovery from medicinal plants has played an important role in the treatment of cancer and most clinical applications of plant secondary metabolites and their derivatives over the half century have been applied towards combating cancer. Of all available anticancer drugs from 1940 to 2002, 40% were natural products or natural product derived, with another 8% considered natural product mimics. Plants have a long history of use in the treatment of cancer, though many of the claims for the efficacy of such treatments should be viewed with some skepticism because cancer, as a specific disease entity, is likely to be poorly defined in terms of folklore and traditional medicine. According to [9], over 50 % of the drugs in clinical trials for anticancer properties were isolated from natural sources or are related to them. Several natural products of plant origin have potential value as chemotherapeutic agents. Some of the currently used anticancer agents derived from plants are podophyllotoxin, taxol, vincristine and camptothecin [10]. The areas of cancer and infectious diseases have a leading position in utilization of medicinal plants as a source of drug discovery. Among FDA approved anticancer and anti-infectious drugs, drugs from natural origin have a share of 60 % and 75 % respectively [11], yet there are still a number of plants that have an anticancer potential but have not been fully investigated. The National Cancer Institute collected about 35,000 plant samples from 20 countries and have screened around 114,000 extracts for anticancer activity [12]. Over 3000 species of plants with antitumour properties have been reported [13]. Cancer is one of the most prominent diseases in humans and currently there is considerable scientific and commercial interest in the continuing discovery of new anticancer agents from natural product sources [14]. Mortality caused by cancers is increasing throughout the world and it is predicted that more than 13.1 million deaths will occur due to cancer worldwide by 2030 [15]. A large number of chemopreventive and chemotherapeutic agents have been discovered from natural products and they provide a promising strategy to Fight cancer by inducing apoptosis in malignant cells [16,17]. The most important problem in cancer treatment is destroying tumor cells in the presence of natural cells, without damaging natural cells.

It is believed that anticancer effects of plants develop by suppressing cancer’s stimulating enzymes, repairing DNA, stimulating production of antitumor enzymes in cell, increasing body immunity and inducing antioxidant effects [18]. The following are some excellent examples of *vinca* alkaloids, vinblastine and vincristine, isolated from *Catharanthus roseus*, etoposide and teniposide, the semisynthetic derivatives of epipodophyllotoxin, isolated from *Podophyllum* species, the naturally derived taxanes isolated from *Taxus* species and the semisynthetic derivatives of camptothecin, irinotecan, and topotecan, isolated from *Camptotheca acuminata*. [19].

The present study aims to evaluate the anticancer activity of methanolic extracts of *Rotheca serrata* Stane and Mabb. (L.) roots and leaves in both the cancer cell lines (MCF-7 and Neuroblastoma SH-SY5Y).

## MATERIALS AND METHODS

### 1. Plant material

The plant material of *Rotheca serrata* were collected from Panhala, Kolhapur district, Maharashtra, India. The herbaria were prepared and deposited in acronym SUK. It was authenticated by Dr. Lekhak SUK Department of Botany Shivaji University, Kolhapur.

#### 1.1. Preparation of Plant extracts

Air dried plant material of *Rotheca serrata* (root and leaf) was coarsely powdered and used preparation for 10% methanolic extracts. The extracts were passed through Bacterial filter which served as stock solution and used for further investigation

### 2. Preparation of culture media for cell line culture

Dulbecco’s Modified Eagle Medium with High Glucose (DMEM-HG) supplemented with 10% Foetal Bovine Serum (FBS), 1% L-glutamine, 1% penicillin-streptomycin antibiotic solution and was used for culturing MCF-7-Human Breast cancer cell line (Catalogue - ATCC^®^ HTB-22™). Minimum essential Medium (MEM)+F12 medium supplemented with 10% Foetal Bovine Serum (FBS), 1% L-glutamine, 1% penicillin-streptomycin antibiotic solution was used to culture SH-SY5Y-Homo sapiens bone marrow neuroblast. (Catalogue - ATCC^®^ CRL-2266™).

1X Dulbecco’s Phosphate Buffered Saline (DPBS), 0.25% Trypsin-EDTA solution, MTT reagent, Dimethyl Sulfoxide (DMSO), antibiotics, media etc. were all purchased from HiMedia, India.

### 3. Preparation cell for MTT assay

MCF-7 and SH-SY5Y cells were cultured and maintained on respective media. Cells were suspended at 2,50,000 cells/flask in a total volume of 10 ml. All the cells were trypsinized with 1% L-glutamine and 1% penicillin-streptomycin antibiotic solution. The cells were pipetted in 96 well plates at the rate of 1.0 × 10^4^ cells/0.1 ml. The cell cultures were maintained in a 5% CO_2_ incubator at 37^0^C. MTT assay (3-(4,5-Dimethylthiazol-2-yl)-2, 5-diphenyltetrazolium bromide) assay method carried out by Lee [20] was followed accordingly.

1. The trypsinized cells were aspirated into 5ml centrifuge tubes. Cell pellet was obtained by centrifugation at 300 xg. The cell count was adjusted, using DMEM HG medium, such that 200μl of suspension contained approximately 10,000 cells.
2. To each well of the 96 well microtitre plate, 200μl of the cell suspension was added and the plate was incubated at 37°C and 5% CO_2_ for 24 h.
3. After 24 h, the medium was aspirated. 200μl of different test concentrations (25, 50, 100, 200 and 400μg/ml) of plant extract (*R. serrata* roots and leaves) test drugs were added to the respective wells. The wells devoid of plant extract served as negative control. The plate was then incubated at 37°C and 5% CO_2_ for 24 h.
4. The plate was removed from the incubator and the drug containing media was aspirated. 200μl of medium containing10% MTT reagent was then added to each well to get a final concentration of 0.5mg/ml and the plate was incubated at 37°C and 5% CO_2_ for 3 h.
5. The culture medium was removed completely without disturbing the crystals formed. Then 100μl of solubilization solution DMSO was added and the plate was gently shaken in a gyratory shaker to solubilize the formed formazan.
6. The absorbance was measured using a microplate reader at wavelengths 570 nm. The percentage growth inhibition was calculated as follows: **% Growth inhibition of cells = 100 - (Test OD/non treated OD)* 100 exposed to treatment**
7. Concentration of test drug required to inhibit cell growth by 50% (IC_50_) was then generated from the dose-response curve for the respective cell line.
8. The assay measures the cell proliferation rate and conversely, when metabolic events lead to apoptosis or necrosis, the reduction in cell viability. The same procedure was followed for SH-SY5Y using culture medium MEM.

### Statistical analysis

The results of the data were expressed as the mean ± standard error of 6 independent determinations in two separate experiments. Statistical data was performed by using one-way ANOVA while Significance of result is calculated from p value.

## RESULTS

The spreading of cancer is increasing globally and the percentage of deaths caused by this lethal disease is rising particularly in the developing countries. Scientists and researchers have turned to affordable herbal medicine in treatment of complicated diseases like cancer. This is because treatments of patients with chemical therapy have serious side effects and cost of treatment is too high. Recently, herbal medicines are coming to play a more vital role in the reduction and prevention of cancer. Plants like *Nigella sativa, Acacia seyal, Allium sativum, Olea europaea* and *Vitis vinifera* have been studied in for immunomodulatory and cancer treatment purposes. These studies have resulted in isolation of principle compounds with promising cancer treating therapy. Resveratrol is a leading example isolated from *V. vinifera* applied effectively in treating cancer [21].

The efficacy of *Clerodendron serratum* against Dalton’s Ascitic Lymphoma has been reported in previous studies [22]. Similarly, studies done on *in vivo* anticancer activity of (*Rotheca serrata) Clerodendrum serratum* roots on Dalton’s lymphoma ascites has confirmed that methanolic extract of *Clerodendrum serratum* exhibits anticancer activity at the dose 100 mg and 200 mg/kg body weight [23].

In present investigation cancer cell lines MCF-7 and SH-SY5Y were used to study anticancer activity of *Rotheca serrata* instead of direct animal studies on Dalton’s lymphoma ascites in albino mice.

### MTT mechanism

The reduction of tetrazolium salts is widely believed as a reliable way to study cell proliferation.

The effect of test extract on the cellular proliferation and viability were determined using 3- (4,5Dimethylthiazol-2-yl)-2,5-diphenyltetrazolium bromide (MTT) assay method [20]. Reduction of tetrazolium by dehydrogenase enzymes is character of metabolically active cell that produces NADH and NADPH. The formazan product (purple crystals) has low aqueous solubility (Fig. 1). Formazan dissolved in dimethyl sulfoxide (DMSO) was quantified and the intensity of the product colour was measured at 570 nm. Which was directly proportional to the number of living cells in the culture [24].

**Fig.1.**
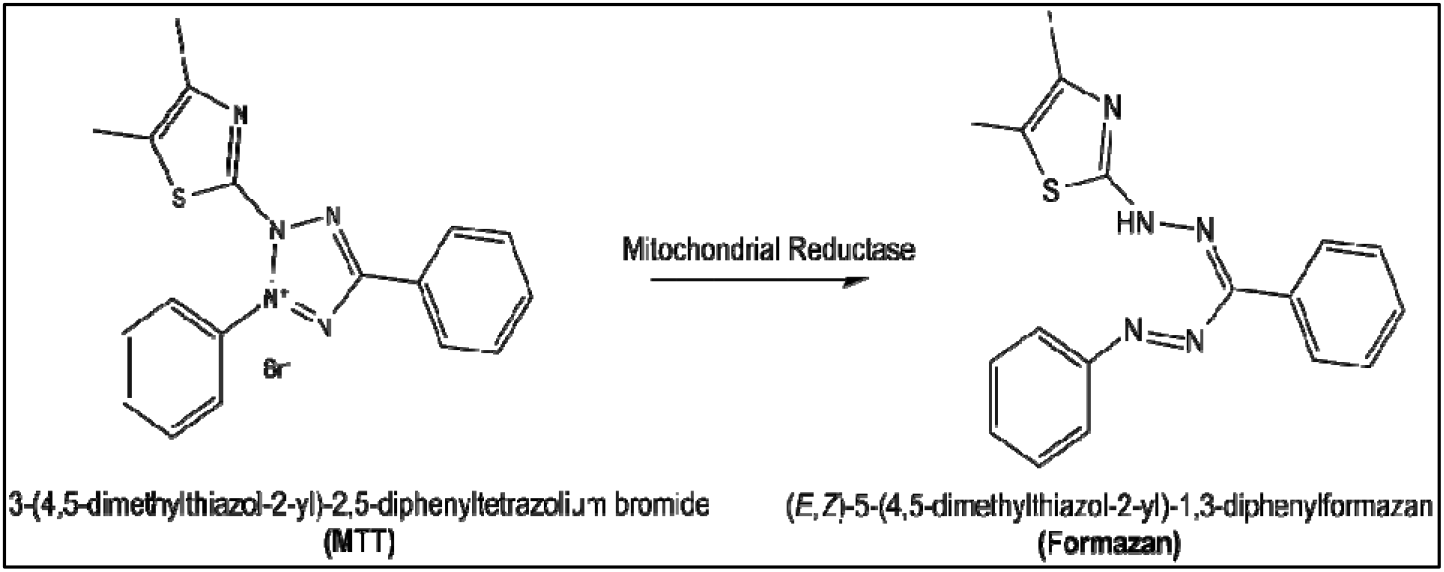
MTT mechanism.

Untreated cells (basal) were used as control of viability (100 %) and the results were expressed as % viability (log) relative to control.

The anticancer activity (cytotoxic effect) was tested for human breast cancer cell line (MCF −7) and neuroblastoma cell line (SH-SY5Y). The concentrations made for evaluation of cytotoxicity were 25 μg/ml, 50 μg/ml,100 μg/ml, 200 μg/ml and 400 μg/ml for MCF – 7 cell lines and SHSY5Y cell lines. Each extract was evaluated in triplicates by serial dilution method.

IC 50 values of test extracts of *Rotheca serrata* (root and leaf both) causing 50% cell death are depicted in Table 1. Among the concentrations used for evaluation of cytotoxicity for breast cancer (MCF – 7) cell lines 400 μg /ml of methanolic extract of root sample was most effective in growth inhibition of cancerous cells. Similar results were obtained for methanolic leaf extracts, where 400 μg /ml was found to be best in inhibiting the growth of cancerous cells. The root methanolic extract showed 5.14 % viability of cancerous cells (Fig. 2) with morphological changes observed in cancerous cells. These changes were mostly related to bursting of cells after treatment (Figs. 3b–3f) was given as compared to control (Fig. 3a). While the leaf methanolic extract showed 8.1 % viability (Fig. 4) with morphological changes shown in (Figs. 5a–5f). Among leaf and root extracts, the IC 50 value of root was calculated as 61.8259 ± 7.428 μg/ml with p-value showing 0.006013 which is significant at p < 0.05. While for leaf IC 50 value was 78.1497 ± 6.316μg/ml with p value 0.0062, which is significant at p < 0.05.

**Fig. 2.**
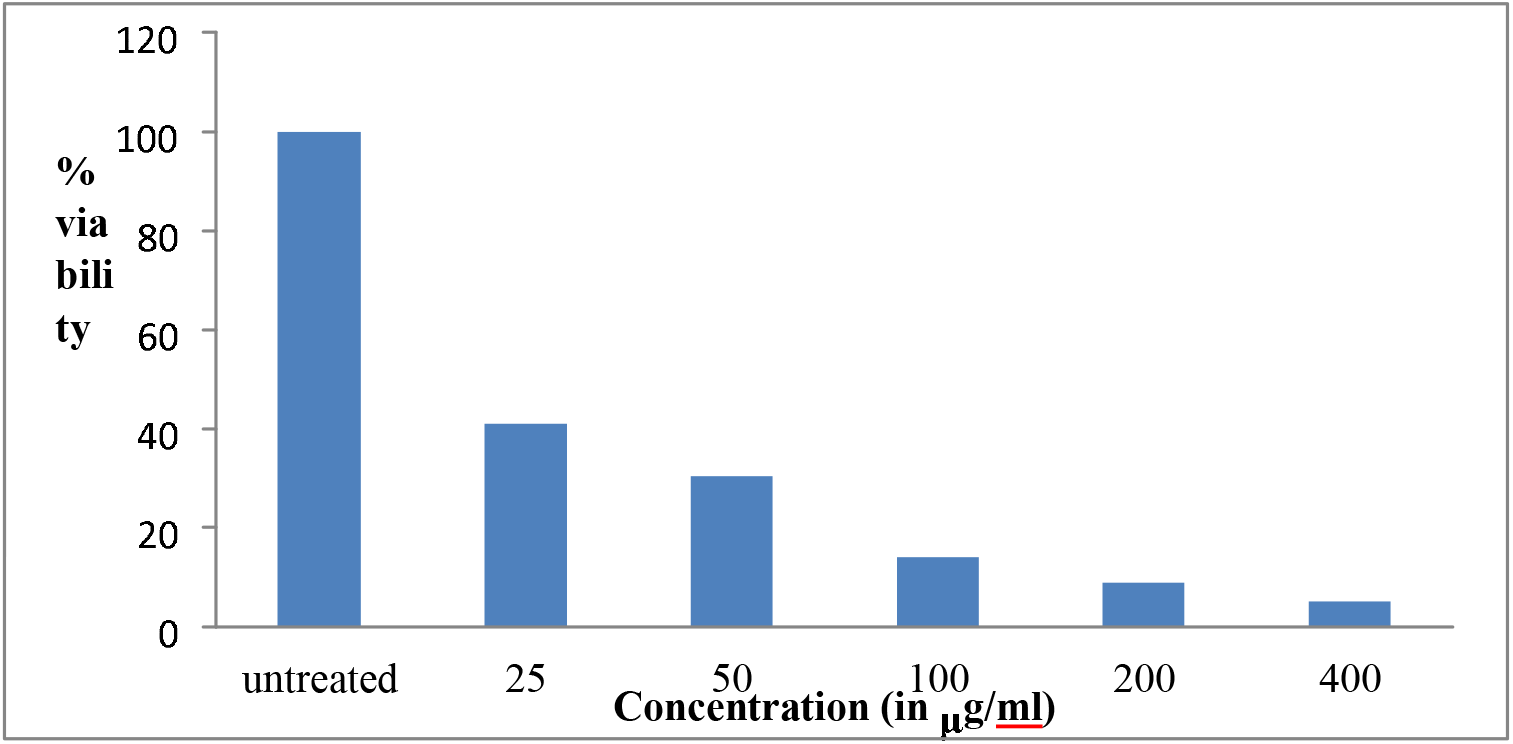
Percent viability of MCF-7 cell line treated with root extract of *R. serrata*.

**Fig. 3.**
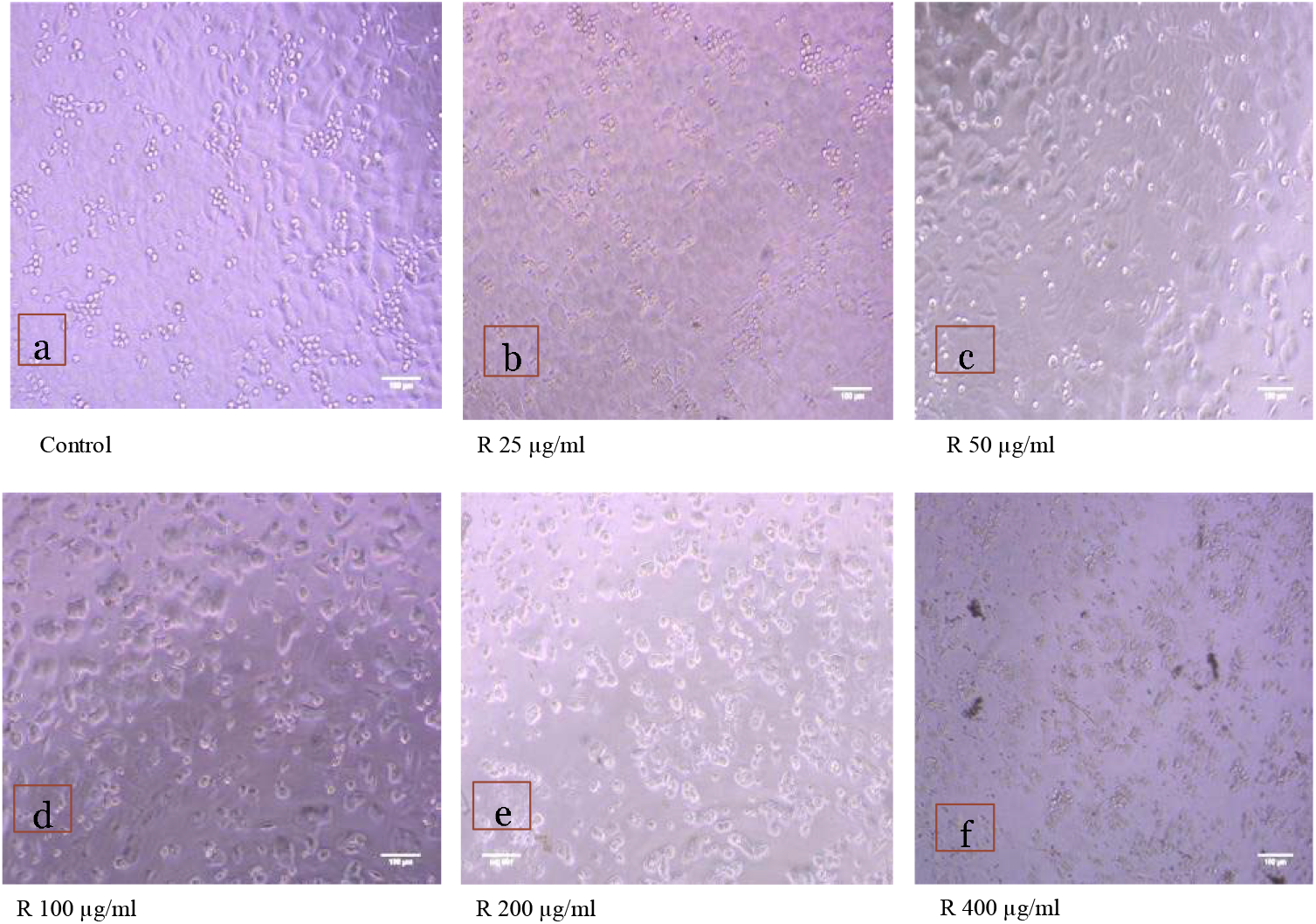
**a-f** Morphological changes showing inhibition of MCF-7 breast cancer cell lines treated with root extract.

**Fig. 4.**
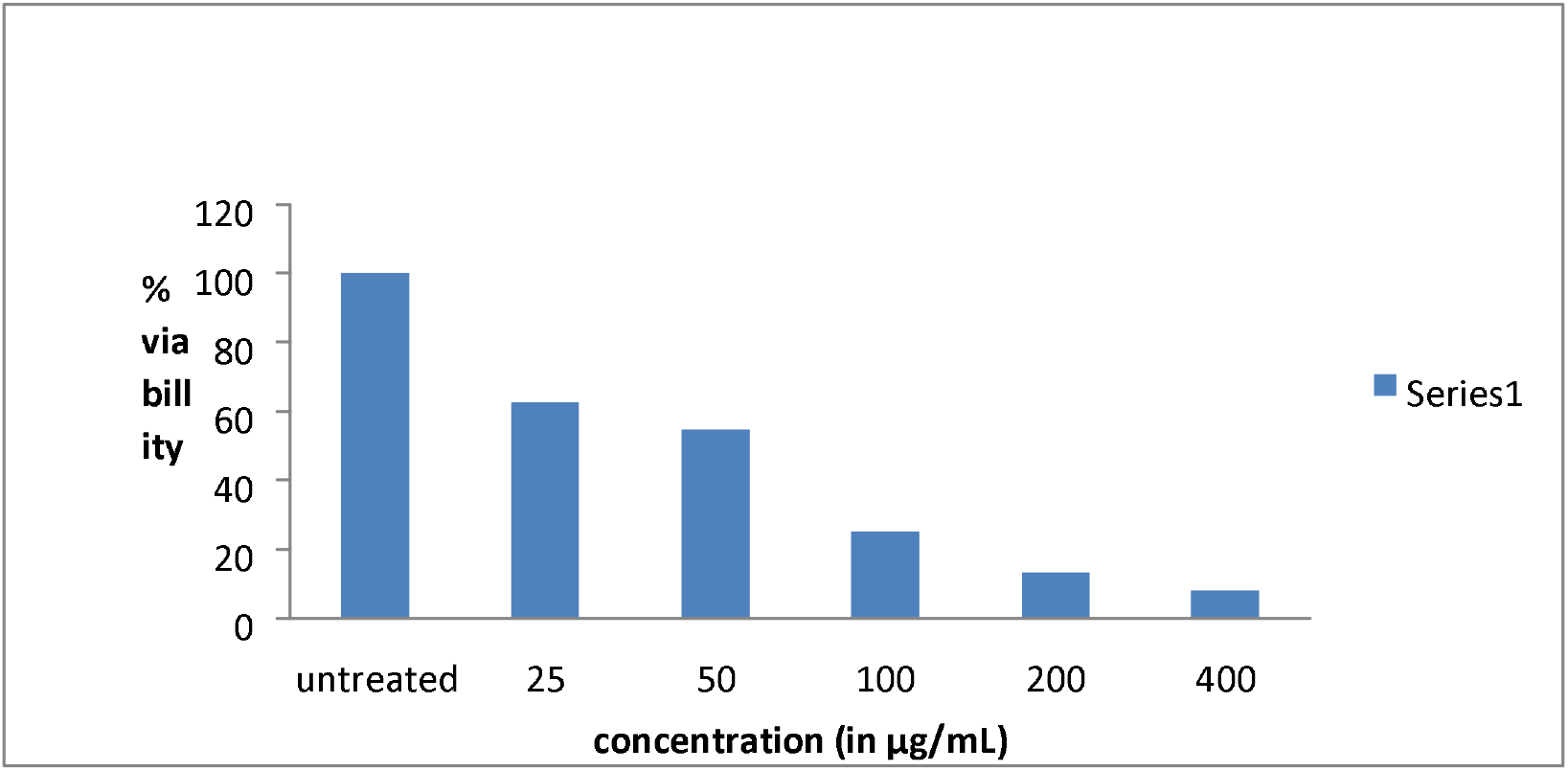
Percent viability of MCF-7 cell line treated with leaf extract.

**Fig. 5.**
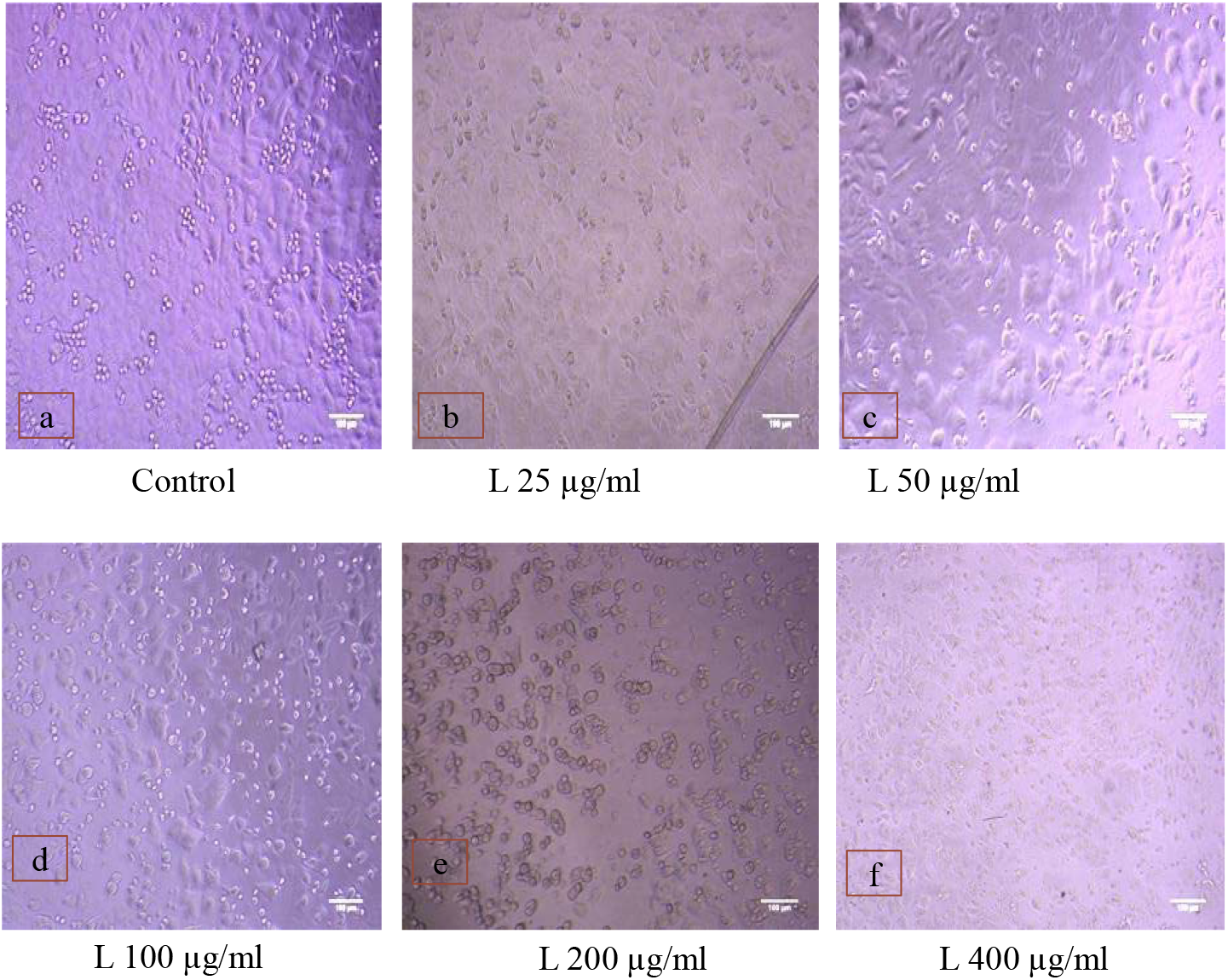
**a-f** Morphological changes showing inhibition of MCF-7 breast cancer cell lines treated with leaf extract.

**Table 1.**
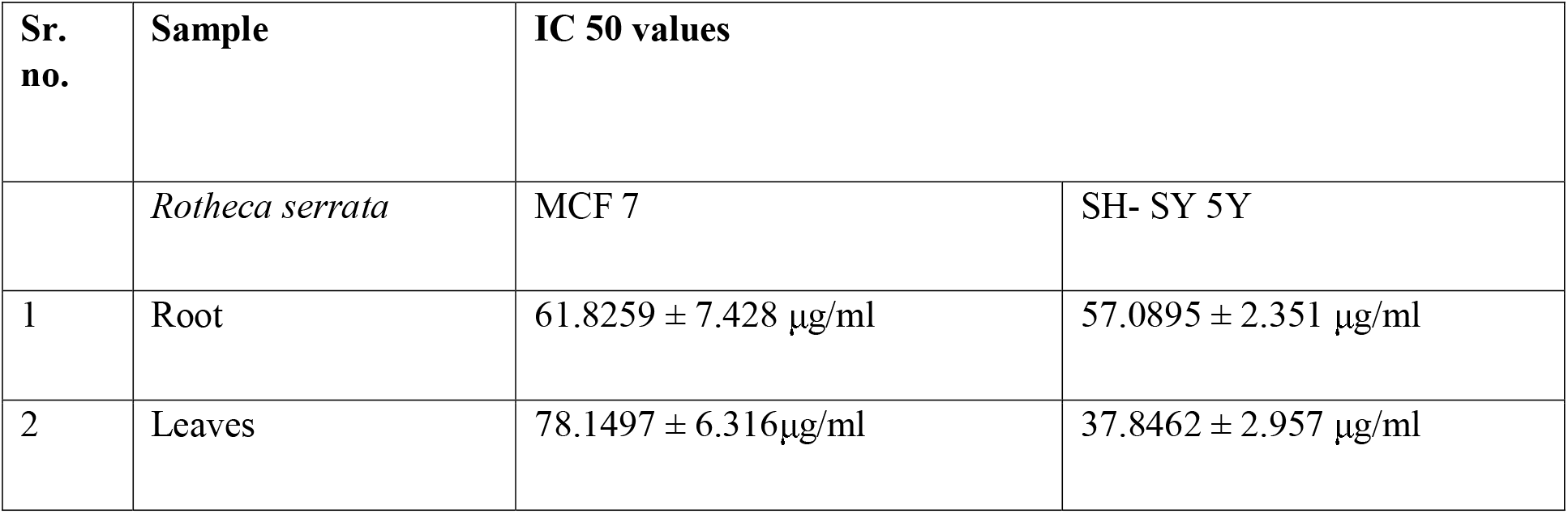
IC50 of *Rotheca serrata* root and leaf extracts on human cancer cell lines (MCF-7 and SH-SY5Y)

In case of neuroblastoma (SH-SY5Y) cell lines the 400μg/ml concentration of leaf methanolic extract showed effective inhibition of cancer cells with IC 50 value 37.8462 ± 2.957 μg/ml with p-value is < 0.00001. The result is significant at p <0.05. The root methanolic extract showed IC 50 value 57.0895 ± 2.351 μg/ml. The p-value is 0.249871 due to which result is *not* significant at p > 0.05 in case of root extract. The root and leaf extract of *R. serrata* showed 7.05 % and 13.97 % viability of neuroblastoma SH-SY5Y cancerous cells which were depicted in Figs. 6 and 8 respectively. The post treatment cell bursting is shown in Figs.7 and 9 respectively.

**Fig. 6.**
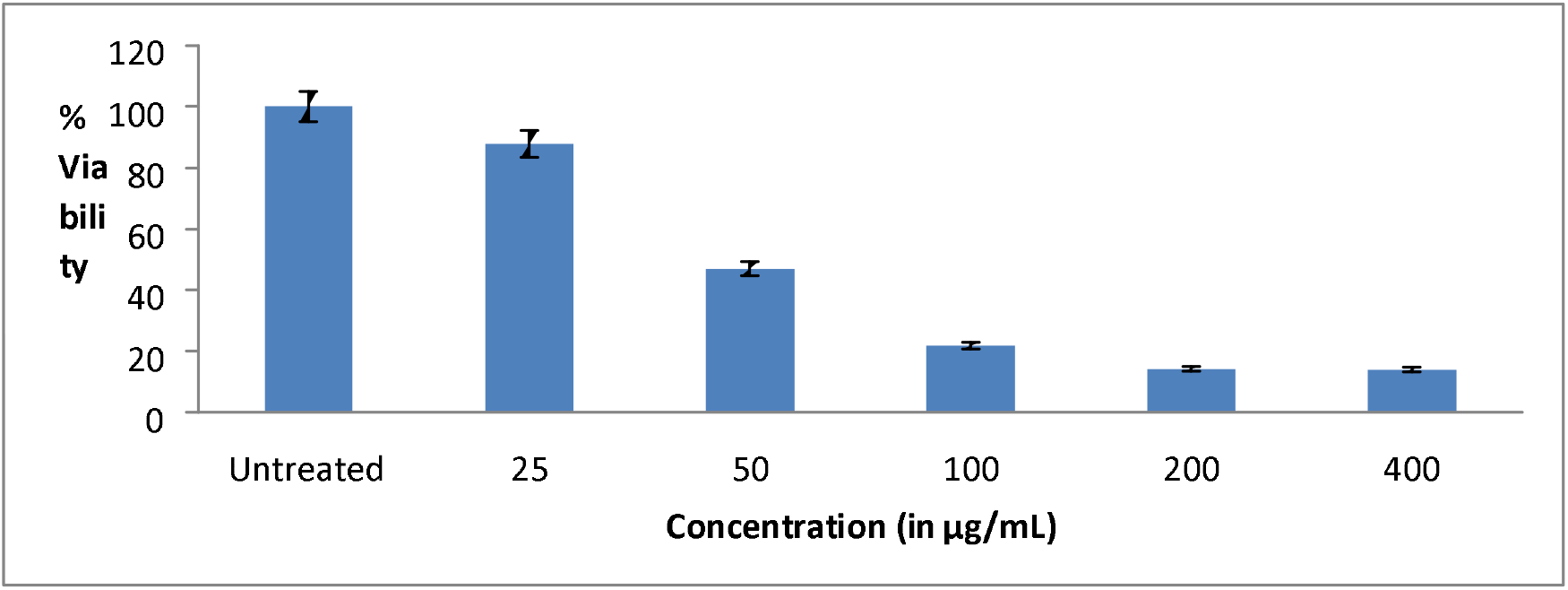
Percent viability of SHSY-5Y cell line treated with root extract of *R. Serrata*.

**Fig. 7.**
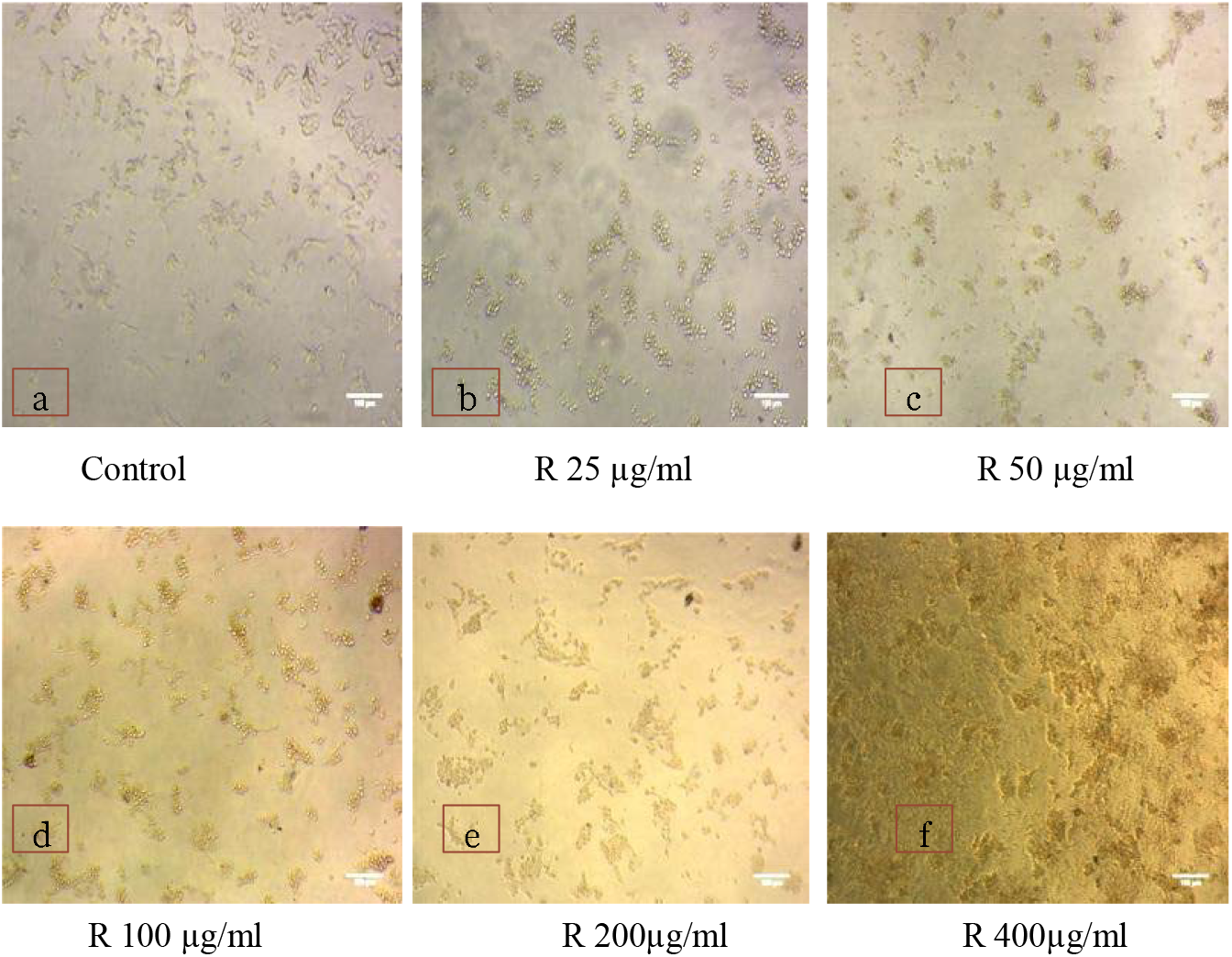
**a-f** Morphological changes showing inhibition of SHSY-5Y neuroblastoma cell lines.

**Fig. 8.**
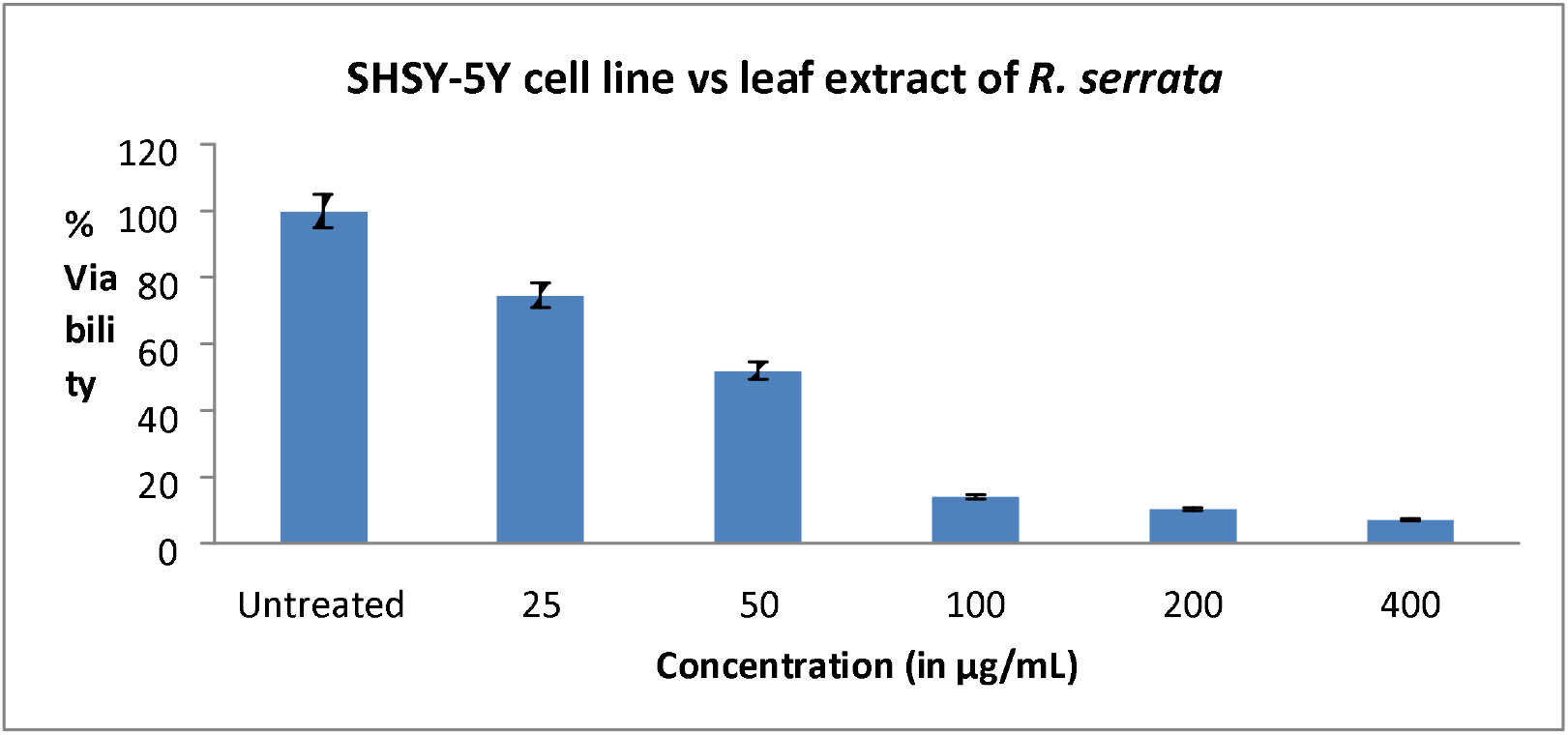
Percent viability of SHSY-5Y cell line treated with leaf extract of *R. serrata*.

**Fig. 9.**
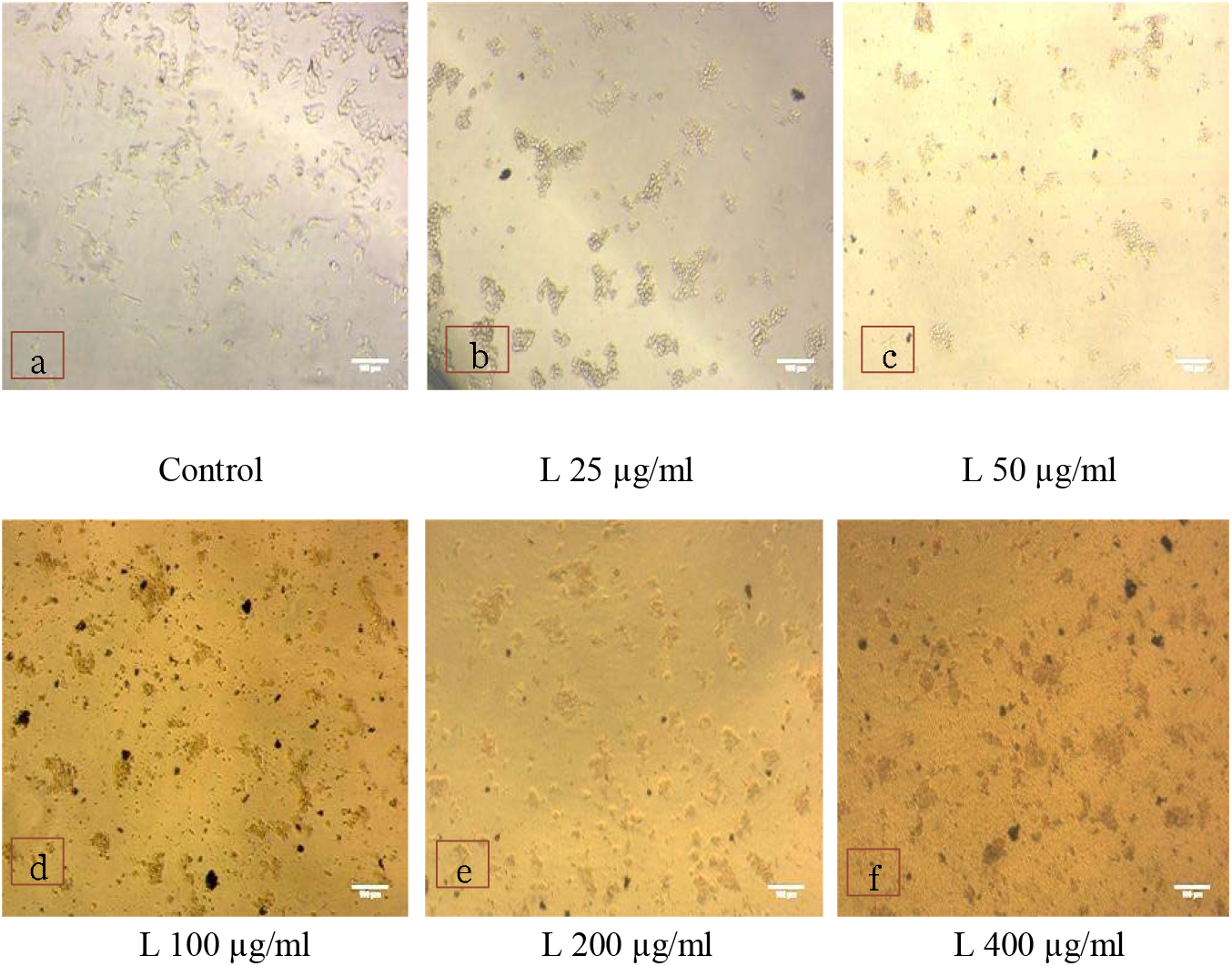
**a-f** Morphological changes showing inhibition of SHSY-5Y neuroblastoma cell lines.

The results showed that methanolic extract of root of *Rotheca serrata* IC 50 (61.8259 ± 7.428 μg/ml) for MCF - 7 cell lines. While IC 50 (37.8462 ± 2.957 μg/ml) for SH-SY5Y cancer cell line in methanolic extract of leaf of *Rotheca serrata* (Table 1). These studies revealed that methanolic extracts of *Rotheca serrata* have high anticancer potential. The results also confirmed that anticancer effect is concentration dependent. As the concentration increases, the anticancer activity (Cytotoxicity level) was found to increase this trend was followed till IC 50 was achieved.

The presence of secondary metabolites effectively inhibits the growth of cancerous cells. According to [25]. plant sample with IC 50<50μg/ml considered to be highest anticancer property while those having IC50<100μg/ml considered as strong anticancer property. Thus, the results are in concurrence with those shown in *Bougainvillea spectabilis, Equisetum hymela, Chenopodium ambrosoides* etc. [25]. The results will be helpful for pharmaceuticals to develop the drug from plant resource which is ecofriendly to treat the cancer patients.

Medicinal compounds like Quercetin, isorhamnetin, kamfrolinalol, alphapinin, limonene and myrecene of *Artemisia absinthium* are been studied for their anticancer activity. Quercetin has shown to inhibit growth of many cancer cells such as MCF-7 and isorhamnetin inhibits growth of MB-435, SKMEL-5, Du-145, MCF-7 and DLD [26]. The cytotoxic effects of essential oil *Rosa damascenes* was reported on lung cancer cell lines (A549) and breast cancer cell lines (MCF7). The ethanolic extract of the *Rosa damascenes* was shown to have inhibitory effect on cervical cancer cells (HeLa) [27]. Glycyrrhizin, a triterpene glycoside of *Glycyrrhiza glabra* is the main compound in root which acts as an anti-proliferative agent against tumor cells, especially breast cancer cell lines MCF-7 and HEP-2. It plays its role by inducing apoptosis [28,29]. *Thymus vulgaris* inhibits growth of human breast cancer and colorectal cancer this is due to presence of phytochemicals like thymol and carvacrol [30].

In similar studies, *Nardostachys jatamansi* was reported to have 54% and 91% inhibition against neuroblastoma cell line [31]. Similarly, *Saussurea lappa* root extract was found to induce dose dependent activity against neuroblastoma cell line [32]. In the results obtained for evaluation of cytotoxic effect of leaf extract of *Citrus limon* on neuroblastoma SH-SY 5Y cell lines showed IC 50 value 72.52 μg/ml [2] with significant dose dependent inhibition of SH SY5Y cell lines while in present findings the leaf extract of *Rotheca serrata* showed IC 50 value 37.8462 ± 2.957 μg/ml which confirms that higher anticancer activity profile as compared to *Citrus limon*.

Methanolic extract of *Hiptage benghalensis* exhibited higher inhibitory effect against tumor cells of HeLa, MCF-7, and IMR-32 cells, with IC 50 values 50.73, 47.90 and 53.76 μg/ml respectively [33]. The anticancer activity of *Leea indica* has shown that methanolic extract significantly inhibited the growth of DU-145 cell line with IC 50 value 529.44±42.07 μg/ml and PC-3 cell line with IC 50 value 547.55±33.52 [34]. In other study 100μg/ml aqueous and methanolic extract of *M. nigra* inhibited 89.5-31.99% of HeLa cell line [35].

## Conclusion

It can be concluded that *R. serrata* possess anticancer activity against breast cancer cell line (MCF-7) and neuroblastoma (SH-SY 5Y) cell lines. This study may help in opening new avenue for cancer research as well as plant-based cancer treatments.

## Supporting information

Abstract

cover letter

structured abstract

## Notes

### Competing Interest Statement

The author JG is thankful to the Department of Botany, Shivaji University and University Grants Commission (UGC), New Delhi, INDIA for providing CSIR UGC NET JRF fellowship for the research work.

### Summary of Updates

All corrections suggested are rectified.

