## Supplementary material for "Cytotoxic effect of *Rotheca serrata* on cancer cell lines MCF-7 and Neuroblastoma SH-SY5Y": structured abstract

**INTRODUCTION:**

Although surgery, chemotherapy, radiation, cryoablation, immunotherapy, targeted drug therapy, hormone use etc. are used for treatment of breast and neuroblastoma, plant-based drugs need to be studied extensively to mitigate the complications encountered due to higher dose regimen.

**AIM OF THE STUDY:**

This work aimed to study the anticancerous effect of *Rotheca serrata* (root and leaf) on cancer cell lines MCF-7 (breast cancer cell line) and Neuroblastoma SH-SY5Y.

**MATERIALS AND METHODS:**

This investigation was a preliminary one which supported the retrospective and safe use of plants as described in Ayurveda. Dulbecco's Modified Eagle Medium with High Glucose (DMEM-HG) for culturing MCF-7- Human Breast cancer cell line and Minimum essential Medium (MEM)+F12 medium for culturing SH-SY5Y- Homo sapiens bone marrow neuroblast. Were used. MTT assay measured the cell proliferation rate and conversely, when metabolic events lead to apoptosis or necrosis, the reduction in cell viability.

**RESULTS:**

The results indicated that the Methanolic extract of *Rotheca serrata* (root and leaf) showed high anticancer activity. Different concentrations of plant extracts (25, 50, 100, 200, 400 µg/ml) were used to study the anticancerous activity, amongst which the significant results were obtained for 400 µg/ml concentration (both root & leaf). Effective anticancer activity against MCF – 7 breast cancer cells was shown in methanolic extracts and were expressed as IC 50 values; in root (IC 50 value= 61.8259 ± 7.428 µg/ml) and in leaf (IC 50 value = 78.1497 ± 6.316µg/ml). The MTT assay in case of neuroblastoma (SH-SY5Y) cell lines revealed that 400µg/ml concentration of leaf methanolic extract showed effective inhibition of cancer cells with IC 50 value 37.8462 ± 2.957 µg/ml as compared to IC 50 value of root methanolic extract which was 57.0895 ± 2.351 µg/ml.

### CONCLUSION:

*R. serrata* possess anticancer activity against breast cancer cell line (MCF-7) and neuroblastoma (SH-SY 5Y) cell lines. This study may to design plant-based drugs without side effects. Dosage compensation for specific type of cancer needs to be monitored in patients with 1st stage.
